# TRPM7 residue S1269 mediates cAMP dependence of Ca^2+^ influx

**DOI:** 10.1101/269688

**Authors:** Jorrit Broertjes, Jeffrey Klarenbeek, Yasmin Habani, Michiel Langeslag, Kees Jalink

**Affiliations:** Division of Cell Biology I, The Netherlands Cancer Institute, 1066 CX Amsterdam, The Netherlands; Department of Physiology and Biomedical Physics, Medical University of Innsbruck, A-6020 Innsbruck, Austria

**Keywords:** Ca^2+^ signaling, TRPM7, cAMP, PKA, fluorometry

## Abstract

The nonspecific divalent cation channel TRPM7 (transient receptor potential-melastatin-like 7) is involved in many Ca^2+^ and Mg^2+^-dependent cellular processes, including survival, proliferation and migration. TRPM7 expression predicts metastasis and recurrence in breast cancer and several other cancers. In cultured cells, it can induce an invasive phenotype by promoting Ca^2+^-mediated epithelial-mesenchymal transition. We previously showed that in neuroblastoma cells that overexpress TRPM7 moderately, stimulation with Ca^2+^-mobilizing agonists leads to a characteristic sustained influx of Ca^2+^. Here we report that sustained influx through TRPM7 is abruptly abrogated by elevating intracellular levels of cAMP. Using pharmacological inhibitors and overexpression studies we show that this blockage is mediated by the cAMP effector Protein Kinase A (PKA). Mutational analysis demonstrates that the Serine residue S1269, which is present proximal to the coiled-coil domain within the protein c-terminus, is responsible for sensitivity to cAMP.

## 1 Introduction

TRPM7 is a ubiquitously expressed channel-kinase that regulates ion levels, affects gene expression and phosphorylates several target proteins. This versatile protein is involved in many cellular processes including cell survival, proliferation and migration [1,2]. TRPM7 is also involved in other clinically relevant pathological processes, such as anoxic neuronal death [3,4] and cardiac pathology [5,6].

The involvement of TRPM7 in cancer development is increasingly recognized. Breast cancer patients with a high TRPM7 expression have a poor prognosis [7–9] and TRPM7 single nucleotide polymorphisms (SNPs) are associated with breast cancer [10]. Ca^2+^ signals mediated by TRPM7 are thought to facilitate metastasis by inducing the epithelial-mesenchymal transition (EMT). This gives rise to a more motile and aggressive phenotype. Therefore, TRPM7 may be considered a prometastatic protein and an important player in Ca^2+^ driven dissemination of cancer [11].

The TRPM7 protein shows 6 trans-membrane domains, a pore-forming domain, a coiled-coil domain and an α-kinase domain. TRPM7 forms a tetrameric channel present at the plasma membrane that is permeable to divalent ions including Ca^2+^, Mg^2+^, and Zn^2+^[12]. A role on intracellular vesicles, in particular on Zn^2+^ storage vesicles, has also been demonstrated [13] recently. The TRPM7 kinase domain phosphorylates several target proteins, including Myosin heavy chain, and thereby affects cell adhesion and migration [14][15]. The α-kinase can also be cleaved off and translocate to the nucleus where it regulates gene function epigenetically [16,17].

We focused on the role of TRPM7 at the plasma membrane as a Ca^2+^ entry pathway. Extracellular Ca^2+^ levels are about 4 orders of magnitude higher than intracellular levels, and this steep gradient enables Ca^2+^ to fulfil its important role as intracellular messenger involved in a wide variety of cellular processes, including polarization, adhesion, and migration. TRPM7 is also involved in setting the basal Ca^2+^ concentration as its expression and knockdown affect levels of cytosolic Ca^2+^ [14][18] and calcium “sparks” or “flickers”, respectively [19][20][21]. These short lived local Ca^2+^ peaks are thought to coordinate the direction of cell migration.

The mechanism of TRPM7 channel activation is not yet fully elucidated. TRPM7 activity is influenced by many cellular and environmental cues. PIP_2_ hydrolysis has been reported to close TRPM7 channels [22] but on the other hand, we and others found that PIP_2_ hydrolysis by phospholipase C (PLC) may activate the channel [18,23–25] **I**n addition, Mg^2+^ levels, nucleotide concentration, cAMP levels, pH and reactive oxygen species (ROS) all have been reported to affect TRPM7 channel gating, for reviews see Yee *et al.* 2014 and Visser *et al.* 2014 [1,2]. Note that some of these proposed mechanisms have been heavily debated, perhaps because in different studies, different readout methods were used to quantify TRPM7 activity. For example, in whole-cell patch clamping studies, Mg^2+^-free pipette solutions have been used to evoke large outward rectifying TRPM7 currents that are easily quantified [22,26]. We and others noted that the effects of cell signaling on TRPM7 currents as detected in whole-cell patch clamping do not always mirror the effects of these signals on TRPM7-dependent cell-biological read-outs, including migration, adhesion and the regulation of Ca^2+^ levels. We therefore have been applying non-invasive techniques to study TRPM7 activity. Using fluorescent monitoring of intracellular Ca^2+^ levels we reported that in TRPM7-overexpressing mouse neuroblastoma cells (N1E-115/TRPM7), addition of PLC-activating agonists causes a sustained Ca^2+^ influx that is not observed in N1E-115 control cells [14,18]. This result contrasts with the inhibitory action of PLC activation as detected in whole-cell patch clamp studies [22]. Importantly, activation of TRPM7 by PLC-coupled agonists was confirmed in perforated-patch clamp experiments [18,27,28], in other cell lines [25], and it is in line with reported biological effects downstream of TRPM7 which appear to be enhanced, rather than inhibited, by PLC-activating stimuli [14,19,23,24,29]. Thus, Ca^2+^ fluorometry offers a convenient and highly sensitive readout to study TRPM7 activation at the plasma membrane and avoids potential problems caused by internal perfusion.

Using the N1E-115 neuroblastoma system, we here describe that the TRPM7-mediated sustained Ca^2+^ influx is abrogated upon elevation of cAMP levels. This effect is PKA dependent and involves serine residue S1269 in the carboxyl terminus of TRPM7. Our results reveal a hitherto unrecognized control mechanism for TRPM7 and emphasize the complex cellular regulation of this versatile protein.

## 2 Materials and methods

### 2.1. Cell culture and transfection

Mouse neuroblastoma cells overexpressing TRPM7-WT (N1E-115/TRPM7) have been previously described [14]. The S1269A mutation was introduced in wild type TRPM7 cDNA in a pTracer vector, using the Phusion Site-Directed Mutagenesis Kit and primers, both from Life Technologies (Waltham, MA, USA). Primers: S1269A-forward: TCACACGAGAATTGGCTATTTCCAAACACT, S1269A-reverse: AGTGTTTGGAAATAGCCAATTCTCGTGTGA, S1224A-forward: CTACATAAAAAGAGCATTACAGTCTTTAGA and S1224A-reverse: TCTAATGATTGTAATGATCTTTTTATGTAG. Constructs were checked by sequencing and inserted as an XhoI-Not1 fragment into a LZRS-neomycin resistant retroviral vector and introduced in the parental N1E-115 cells. Cells were selected for neomycin resistance and adhesive properties. Retroviral transduction resulted in stable and moderate TRPM7-S1269A overexpression to levels comparable to those of WT TRPM7 in N1E-115/TRPM7-WT cells. Total mRNA was extracted using the GeneJET RNA Purification Kit (Thermo Fisher) according to manufactures protocol and cDNA was synthesized using SuperScript II rtPCR enzyme (Thermo Fisher). PCR was performed using SYBR-Green (Takara) using the following primers: Fw TAGCCTTTAGCCACTGGACC and Rv GCATCTTCTCCTAGATTGGCAG. Expression levels in N1E-115 wild type cells were set to 1. Cells were cultured in Dulbecco’s Modified Eagle Medium (DMEM) containing 10% Fetal Calf Serum (FCS). Penicillin and streptomycin added at 100 μg/ml each were from Gibco, Life Technologies (Waltham, MA, USA). Cells were seeded on 24-mm glass-coverslips in 6-well plates. Transient transfections were done using 1 μg DNA and 3 μg Polyethylenimine (PEI; Polysciences Inc. Warrington, PA, USA) per well.

### 2.2. Materials and constructs

The PKA regulatory and catalytic subunits were transiently expressed [30]. Fluorometry was done using Förster/Fluorescence Resonance Energy Transfer (FRET) based biosensors for cytosolic Ca^2+^ and cAMP. For Ca^2+^ the troponin C-based sensor, Twitch-2B was used [31]. Additionally, for cAMP the Epac-based sensor Epac was used [32]. The compounds bradykinin, forskolin, IBMX and ionomycin were from Calbiochem-Novabiochem Corp. (La Jolla, USA). Prostaglandin E1 (PGE1) was from Sigma-Aldrich (Zwijndrecht, The Netherlands). The PKA inhibitor H-89 was obtained from (Biolog, Bremen, Germany). The Epac-selective cAMP analogue 007-AM (8-pCPT-2-O-Me-cAMP-AM) was from Cayman Chemical (Michigan, USA). CaCl_2_ salt was from Merck (Darmstadt, Germany). Fluorescent Ca^2+^ indicators Oregon Green 488 BAPTA-1-AM and Fura Red-AM were from Invitrogen, Life Technologies (Waltham, MA, USA).

### 2.3. Fluorometric Ca^2+^ and cAMP measurements

Fluorometry was done on an inverted Nikon widefield microscope. A 63x oil immersion lens was used, in combination with a 34°C stage heater. The microscopy medium was kept at a pH of 7.2, using HBS (HEPES buffered saline). This buffer contained 10 mM glucose, 2 mM CaCl_2_, 5 mM KCl, 140 mM NaCl, 1 mM MgCl_2_ and 10 mM HEPES. The excitation of the Cyan Fluorescent Protein (CFP) was set at 425 nm. CFP and Yellow Fluorescent Protein (YFP) emissions were measured simultaneously; using band-pass filters at 470±20 and 530±25 nm respectively. Ca^2+^ measurements were done using FRET biosensors and chemical dyes. The Ca^2+^ FRET sensor Twitch-2B was used. Also, the Ca^2+^ dyes Oregon Green 488 BAPTA-1-AM and Fura Red-AM, were used. No quantitative differences could be detected between these methods, with respect to the TRPM7 Ca^2+^ signals. Before the experiments the FRET ratio was set at 1.0 and a baseline was recorded. After the Ca^2+^ experiments a calibration was done adding ionomycin (10 μM) and a high dose of Ca^2+^ (CaCL_2_, 10 mM).

### 2.4. Statistics

Each figure is representative of experiments on at least three different days. For clarity, a representative Ca^2+^ trace is shown. The Ca^2+^ peaks were normalized 0 to 1, using feature scaling with the following formula: *Xi* 0 to 1 = (*Xi*-*Xmin*)/(*Xmax*-*Xmin*). Where relevant, the respective traces are complemented with quantification, based on the indicated number of Ca^2+^ traces. Statistical significance was determined using a two-tailed unpaired Student’s t-test. The following convention was used to indicate statistical significance: P ≤ 0.05 = *, P ≤ 0.01 = **, and P ≤ 0.001 = ***. Error bars indicate the Standard Error of the Mean (SEM) for the indicated number of measurements.

### 2.5 PKA consensus site prediction

On three different prediction web servers for PKA phosphorylation sites, S1224 and S1269 were indicated as PKA consensus sites. These servers were: NetPhos3.1 [33], pkaPS [34] and GPS 2.0 [35]

## 3 Results and discussion

### 3.1. TRPM7-mediated sustained Ca^2+^ influx terminates upon cAMP elevation

In N1E-115 control cells addition of bradykinin (BK) evokes a single, brief Ca^2+^ spike that terminates after approximately one minute. We and others previously showed that moderate overexpression of TRPM7 changes the kinetics of the BK response in that the initial transient response is followed by a more sustained phase of elevated Ca^2+^ levels that lasts several minutes (**Fig. 1A**) [14,18,25]. We also showed that the initial transient response is due to IP_3_-mediated release of Ca^2+^ from internal stores, whereas the sustained Ca^2+^ influx phase is strictly dependent on TRPM7 channels functioning at the plasma membrane. Furthermore, sustained Ca^2+^ influx depends on the presence of a Ca^2+^ gradient across the membrane and it is blocked by TRPM7 inhibitors including La^3+^, 2-APB, SKF96365 and Waixenicin-A [18][21]. These results also showed that analysis of calcium levels by ratiometric fluorometry of N1E-115/TRPM7 cells presents a robust and very sensitive readout of TRPM7 activity in intact cells.

When testing a panel of GPCR agonists for induction of sustained Ca^2+^ influx, we unexpectedly observed that pretreatment with prostaglandin E1 (PGE1, 5 μM) prevented the sustained influx following stimulation with BK. While untreated N1E-115/TRPM7 cells show a sustained BK response that lasted for several minutes, in cells pretreated with PGE1 the cytosolic Ca^2+^ returned to baseline within 1 – 2 minutes (**Fig. 1B**).

The calcium-dependent fluorescence levels were quantified at 2 minutes post stimulation with BK. Compared to wild-type N1E-115 cells, in untreated TRPM7 cells Ca^2+^-dependent fluorescence remained significantly elevated (15% +/− 10 %; N= 7; p = 0,006). By contrast, in PGE1 pretreated TRPM7 cells Ca^2+^-dependent fluorescence had returned to near-baseline values after 2 minutes (2% +/− 1%; N = 12; p = 0.265). We next set out to investigate the responsible mechanism. PGE1 fails to activate PLC in these cells [18] but it is well-known to stimulate production of the second messenger cAMP in various cell types. To determine whether a rise in cAMP levels may be involved in inhibition of TRPM7-mediated Ca^2+^ influx, we pretreated cells with forskolin (25 μM) and IBMX (100 μM), which elevates cAMP levels by activating adenylate cyclase and blocking phosphodiesterase, respectively. Indeed, similar to PGE1 pretreatment, in cells pretreated with IBMX + Forskolin (IF) the Ca^2+^ returned to baseline within 1 – 2 minutes (**Fig. 1B**). At 2 minutes post BK addition, Ca^2+^-dependent fluorescence had decreased to 4% +/− 3% (N = 24; p = 0.052) in IF-pretreated cells.

Using our Epac-based FRET biosensor [32] we confirmed that treatment with PGE1 and IF raises cytosolic cAMP levels in N115 cells, as expected (**Fig. 1C**). Strikingly, when added during the sustained phase of calcium entry, addition of IBMX/forskolin abruptly terminated TRPM7-mediated Ca^2+^ influx (**Fig. 1D**). Thus, elevation of cAMP levels abrogates sustained Ca^2+^ influx in N1E-115/TRPM7 cells.

**Fig. 1.**
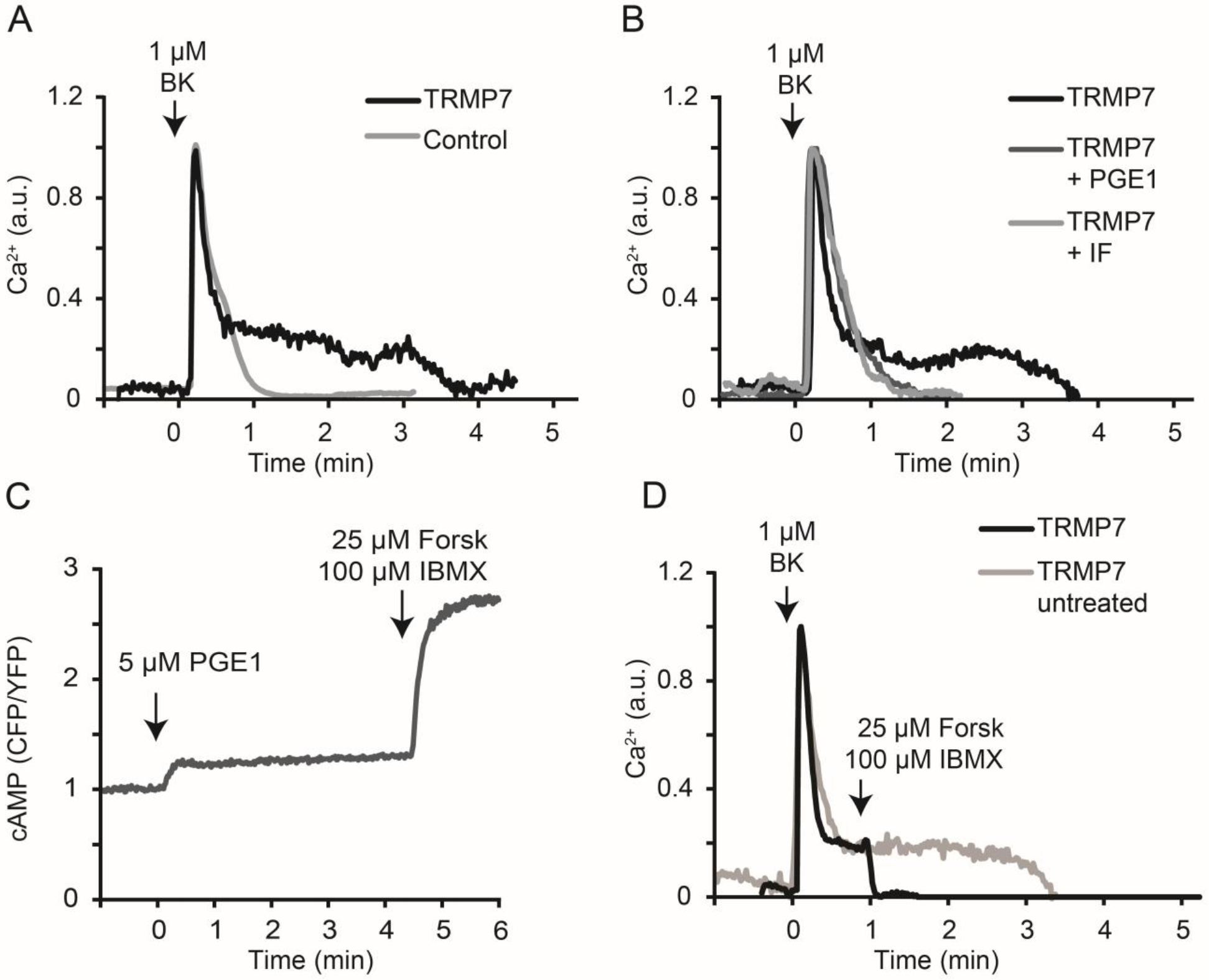
TRPM7-mediated sustained Ca^2+^ influx is abrogated by cAMP elevation. **(A)** Stimulation with bradykinin triggers a sustained influx of extracellular Ca^2+^ in cells overexpressing TRPM7 (black), lasting for several minutes. Control cells (grey) show only a single brief Ca^2+^ transient upon bradykinin stimulation that typically lasts approximately one minute. Calcium levels are expressed as arbitrary units (a.u.). **(B)** Pretreatment with PGE1 (dark grey) or IBMX/forskolin (lighter grey) prevents the sustained influx in TRPM7-WT overexpressing cells. **(C)** Using the Epac-based FRET biosensor it was verified that both PGE1 and IBMX/forskolin treatment elevate cytosolic cAMP concentrations. **(D)** IBMX/forskolin stimulation abruptly terminates the TRPM7-mediated sustained Ca^2+^ influx (arrow). For quantification, see fig. 3.

### 3.2. Involvement of protein kinase A

To determine which cAMP effector protein is involved, we used specific activators and inhibitors for PKA and Epac (Exchange Protein directly Activated by cAMP), the two most prominent targets downstream of cAMP. Initially, we tested for Epac involvement, using the Epac-selective cAMP analogue 007-AM (8-pCPT-2-O-Me-cAMP-AM) [36]. Pretreatment of cells with 1 μM 007-AM had no effect on basal calcium levels and also did not affect the BK-induced sustained influx (**Fig. 2A**). 007-AM did cause rapid FRET changes in our Epac-based biosensor [32], indicating that it readily permeates the membrane and activates the Epac protein (**Fig. 2A**). These experiments exclude a role for Epac in the cAMP-induced termination of Ca^2+^ influx. In contrast, the PKA inhibitor H-89 (10 μM) [37] completely prevented the IBMX/forskolin-induced termination of sustained Ca^2+^ influx (**Fig. 2B**) in 13 out of 13 experiments (p < 0.001). These data indicate that PKA may play an important role in the termination of Ca^2+^ influx through TRPM7.

At rest, PKA is a tetramer of two identical kinase subunits and two regulatory subunits that inhibit activity of the kinases. cAMP binding to the regulatory subunits causes the complex to dissociate, releasing the catalytic subunits from inhibition and allowing them to phosphorylate consensus sequences in a variety of cellular proteins. To further investigate the link between PKA activation and termination of Ca^2+^ influx through TRPM7 we overexpressed PKA subunits individually. It may be expected that overexpression of the catalytic subunit increases PKA activity and therefore reduces influx, whereas overexpression of the regulatory subunit would prevent kinase activity. Note that for these experiments, PKA subunits were overexpressed transiently, as it is difficult to achieve stable overexpression of either catalytic or regulatory subunits of PKA due to its growth-regulatory effects. For these series of experiments, a genetically encoded Ca^2+^ FRET sensor was cotransfected to serve both as a transfection marker and for Ca^2+^ readout. Indeed, overexpression of either PKA-Cat or PKA-Reg significantly affected the duration of the sustained Ca^2+^ influx. The sustained phase in control N1E-115/TRPM7 cells lasted on average 294 seconds (N=5 experiments with 2-3 cells each). Overexpressing the catalytic subunit reduced the influx length to 132 seconds (P = 0.012; N = 4 experiments). Conversely, overexpressing the regulatory subunit elongated the influx to 516 seconds (P = 0.008, N = 3 experiments) (**Fig. 2C**). Taken together, these data strongly indicate that PKA is the effector that mediates termination of the sustained Ca^2+^ influx following elevation of cAMP.

**Fig. 2.**
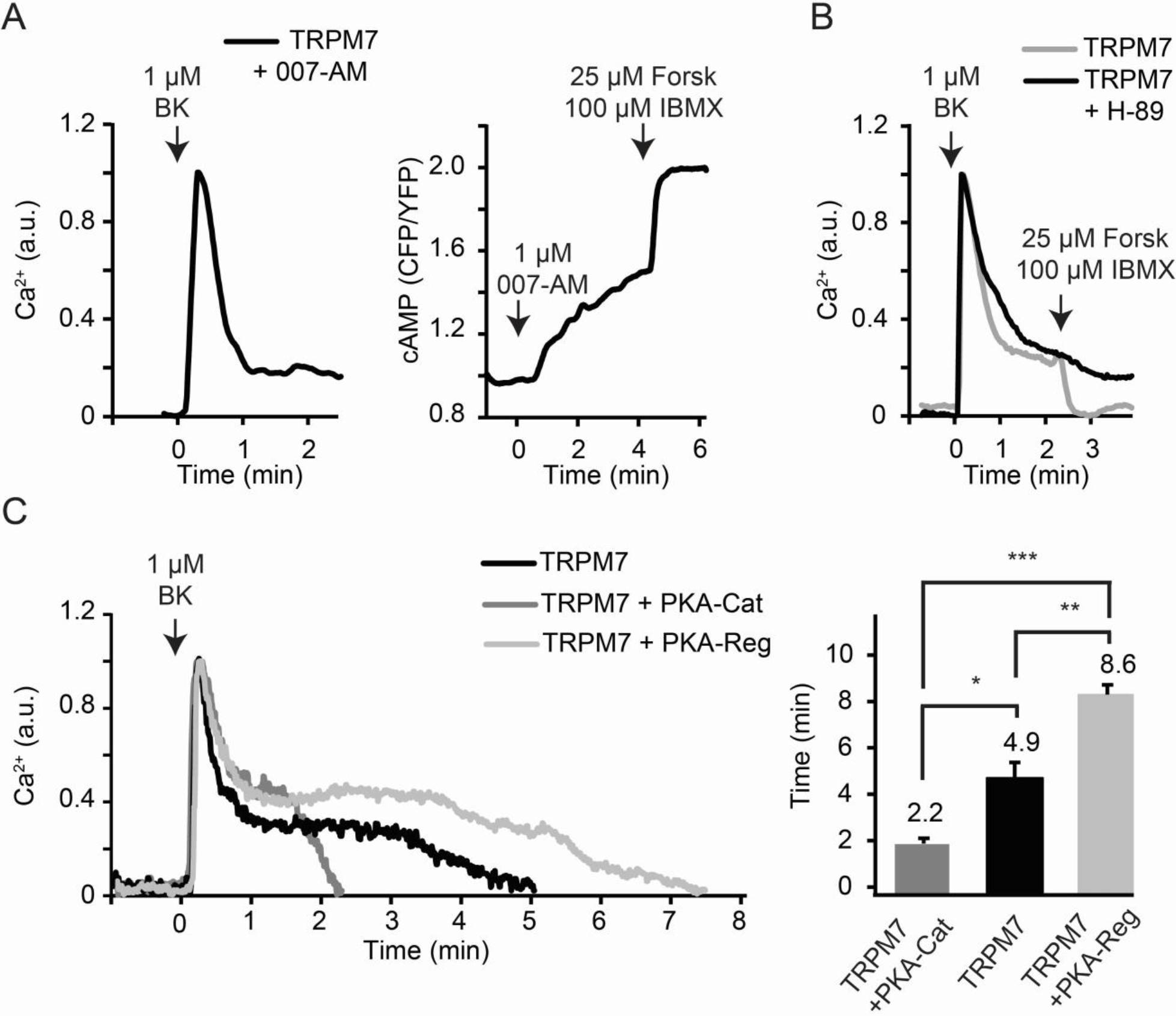
Involvement of PKA in the abrogation of Ca^2+^ influx. **(A)** Left panel, pretreatment with the Epac-selective cAMP analogue 007-AM (1 μM, 5 minutes) did not affect the sustained influx (left panel). Right panel, 007-AM activates the Epac FRET biosensor, verifying its biological activity. **(B)** Pretreatment with PKA inhibitor H-89 (10 μM, 5 minutes) largely blocked sensitivity of Ca^2+^ influx to cAMP, as addition of IBMX/forskolin during the sustained phase was without effect. **(C)** Overexpressing either PKA-Cat or PKA-Reg significantly affected the duration of the sustained Ca^2+^ influx. The left panel shows that the sustained Ca^2+^ influx of TRPM7 alone (black) lasted approximately 5 minutes. Overexpressing the catalytic subunit (dark grey) reduced the duration of the influx to approximately 2 minutes. Conversely, overexpressing the regulatory subunit (light grey) prolonged the influx to approximately 8 minutes. Representative traces from a single experiment are shown; data are quantified in the right panel. *, p < 0.05; **, p<0.01; ***, p<0.001.

### 3.3. A single point mutation, S1269A, renders TRPM7 resistant against cAMP-mediated termination of Ca^2+^ influx

In an attempt to identify possible PKA phosphorylation sites that may mediate cAMP sensitivity of Ca^2+^ influx, we revisited a set of serine point mutants that we had previously prepared for a study into the possible function of the coiled-coil region in TRPM7. Three serines within this set are known to be phosphorylated, namely S1224, S1255 and S1269 [38]. PKA phosphorylates serine (and to a lesser extent threonine) residues of target proteins at PKA consensus sites which consist of arginine residues at positions −3 and often also at −2, and a hydrophobic residue at +1 (RrXSϕ) [39]. Two of the mutants, S1224 and S1269 (see materials and methods) conformed to the PKA consensus signature and consequently we focused on those two for further analysis.

**Fig. 3.**
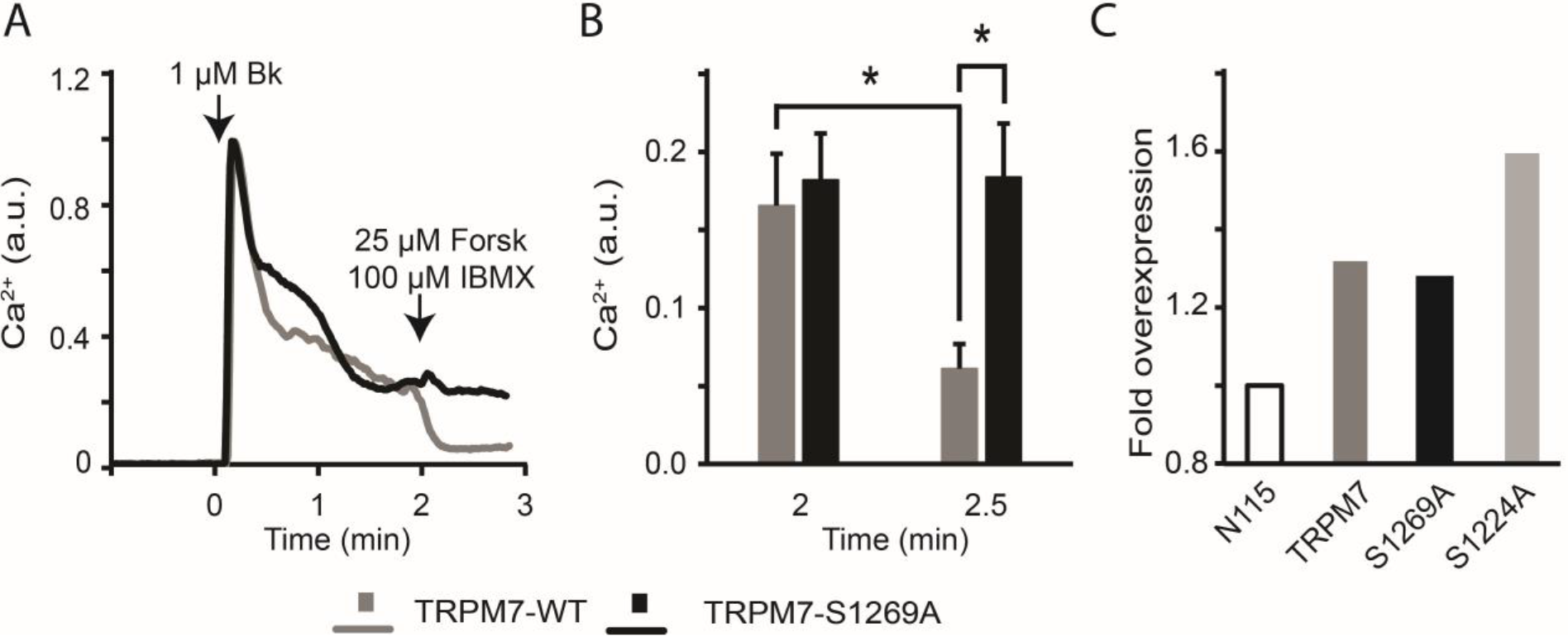
The S1269A phosphorylation-dead mutant was insensitive to cAMP-mediated termination of Ca^2+^ influx. **(A**) Representative traces of Ca2+ responses in TRPM7-S1269A mutant (black) and TRPM7-WT (grey) expressing cells. Note that in the S1269A mutant sustained influx fails to terminate after being challenged with IBMX/forskolin (arrow). **(B)** Quantification of Ca^2+^ levels before (left bars, taken 2 min after addition of BK) and 30 s after (right bars) addition of IBMX/forskolin and expressed as average + SD of N = 7 TRPM7-WT (gray) and N = 10 TRPM7-S1269A (black) cells. IBMX/forskolin were added at 2 minutes post BK addition. In TRPM7-WT expressing cells but not in S1269A mutant expressing cells cAMP caused a significant drop in calcium levels (P = 0.016). The difference between TRPM7-WT and TRPM7-S1269A expressing cells following addition of IBMX/forskolin is also significant (P = 0.013). **(C)** Transduction was checked by qPCR. A moderate 1.3x overexpression of S1269A was seen, as is common for N1E-115-TRPM7 cells. S1224 had a 1.6x overexpression. Note that although moderate, the overexpression levels observed are in line with literature reports [14,40].

N1E-115 cells were retrovirally induced to express either TRPM7-S1224A or TRPM7-S1269A, mutants which cannot be phosphorylated at the respective residues. Proper transduction was checked by QPCR (**Fig. 3C**). Ca^2+^ fluorometry showed that cells expressing TRPM7-S1224A failed to display the characteristic sustained influx of Ca^2+^ seen in TRPM7-WT overexpressing cells when challenged with BK. This may be because S1224A mutants do not localize at the plasma membrane properly, or alternatively, the channel may be defective in activation. In contrast, TRPM7-S1269A expressing cells displayed a prominent sustained Ca^2+^ influx following stimulation with BK. This indicates that the channel was expressed, that it is at the plasma membrane and that it is functional. Strikingly, addition of forskolin and IBMX to those cells failed to terminate the sustained response evoked by stimulation with BK. This implies that S1269A mutant channels have lost their sensitivity for regulation by cAMP (Fig. 3). Thus we conclude that PKA-dependent phosphorylation of S1269 must be a key determinant of agonist-induced Ca^2+^ influx through TRPM7 in these cells.

## Concluding remarks

We found that in cells that overexpress TRPM7, bradykinin stimulation results in a sustained influx of extracellular Ca^2+^ that is sensitive to elevated intracellular cAMP levels. Using pharmacological inhibitors (H-89 and 007-AM) and overexpression of PKA subunits we showed that this effect is mediated by the cAMP effector protein kinase A. We also identified S1269, which is proximal to the coiled-coil region of TRPM7, as a key residue mediating this response. Together, these findings reveal a novel level of complexity in the regulation of TRPM7.

S1269 is found within a PKA consensus phosphorylation site (RELSI) present approximately 15 residues C-terminal from the coiled-coil domain. Three independent online prediction engines identified this site as possible PKA phosphorylation site, but we note that it lacks the preferential (although not obligatory) arginine at position −2 within the consensus motive (RrXSϕ, [39]). This raises the formal possibility that another kinase downstream of PKA is responsible for S1269 phosphorylation. Using mass-spectrometric analysis of tryptic peptides, we attempted to directly demonstrate phosphorylation of TRPM7 in cells transiently overexpressing PKA catalytic subunits, but failed to detect phosphorylation of S1269 or other residues. However, such experiments are challenging and well outside our expertise and therefore we may have simply missed phosphorylation. Kim *et al.* [38] reported that S1269 is subject to phosphorylation but no information on the responsible kinase was included in that study. In a recent study by Cai *et al.* [41] S1269 was also found to be phosphorylated. Interestingly, these authors also showed phosphorylation in a kinase-defective mutant of TRPM7, indicating phosphorylation by an (unidentified) external kinase.

Although TRPM7 is sensitive to many external agents, including receptor agonists, pH, reactive oxygen species and even mechanical stress, the exact mechanisms by which channel gating is regulated are still not fully elucidated. The extensive cytosolic C-terminus of TRPM7 houses phosphorylation sites for several kinases, domains that mediate interaction with PIP_2_, a caspase cleavage site and an α-kinase domain that is important for interaction with PLC [1,2]. Moreover, the protein is heavily autophosphorylated and interacts with several cytoskeletal proteins that, in turn, may convey signals to the channel [40]. It remains to be addressed how S1269 affects agonist-induced Ca^2+^ influx through TRPM7. In TRPM8, Tsuruda *et al.* showed that the C-terminal domain containing the coiled coil is important for tetramer formation [42]. Before embarking on these studies, we had hypothesized that addition of a negatively charged phosphate close to the coiled-coil region could potentially affect channel tetramerization, leading to its inactivation. The second PKA consensus site studied here, S1224, is present within the coiled-coil region, it is target for phosphorylation [38] albeit not in kinase-deficient TRPM7 expressing cells [41]. We showed that expression of S1224A mutant constructs failed to produce channels that mediate discernable Ca^2+^ influx. Nevertheless, more recently a convincing study showed that, at least in Zebrafish, truncated TRPM7 mutants lacking the coiled coil domain form functional channel oligomers [43]. Since our studies did not produce direct experimental support for a role of S1269 phosphorylation in channel oligomerization, alternative hypotheses for the strong phenotype of S1269A have to be considered. Conceivably, cAMP-dependent S1269 phosphorylation could induce an inhibitory conformational change in the protein and/or otherwise affect channel conductive properties. However, given that expression of the mutant phenocopies WT TRPM7 expression in inducing sustained Ca^2+^ influx, we can exclude altered expression or deficient routing to the plasma membrane.

A further complicating factor is that in our study (mutant) proteins are overexpressed in a normal TRPM7-wildtype background. This is because (in line with reported literature) we failed to achieve full TRPM7 knockout in our cells, and only slight overexpression of TRPM7 was tolerated [14]. Up-and downregulation of mRNA levels in TRPM7-dependent malignancies is also typically very modest [44], implicating tight regulation of TRPM7 levels. Consequently, in our study most tetramers of mutant channels can be expected to contain one or more wt TRPM7 proteins which complicates phenotypic characterization. The TRPM7-S1269A mutant cells showed expression levels comparable to those in N1E-115/TRPM7 cells. Importantly, expression of TRPM7-S1269A protein phenocopied N1E-155/TRPM7 WT cells in producing sustained Ca^2+^ influx, except that in these mutants raising cAMP failed to terminate Ca^2+^ influx. The strong and highly reproducible phenotype of S1269A mutant channels indicates that notwithstanding the modest overexpression levels, TRPM7 controls sustained Ca^2+^ influx and S1269 has a key role in mediating cAMP sensitivity of this influx.

At first sight, it might be confusing that the effects of raising cAMP on TRPM7-mediated sustained Ca^2+^ influx appears much faster (i.e., within 10-20 s) than the rise in cAMP (which peaks at 2-3 minutes). Very likely, the culprit is the difference in affinity for cAMP between our FRET sensor (Kd ~ 4 μM) [45] and PKA (Kd of ~230 nM, i.e. ~20-fold) [46]. In addition, unlike the Epac-based FRET sensor, PKA displays strong cooperativity (Hill coefficient >2). Together, This means it will be activated long before the cAMP sensor. On top of that, enzymes (PKA is no exception) often start exerting cellular effects long before they are maximally activated. Finally, we note that the inhibitory effects of cAMP on TRPM7 found in our study contrast with the stimulatory effects documented by Takezawa *et al.* [47]. These authors employed whole-cell patch clamping to show that cells, internally perfused with 100 μM cAMP showed enhanced outwardly rectifying TRPM7 currents in HEK293 cells. Several factors may underlie this discrepancy. First, in whole-cell patch clamp experiments TRPM7 currents are evoked by perfusing the cell interior with Mg^2+^-free internal solution. This triggers large TRPM7 currents that do not necessarily reflect the quite small Ca^2+^ currents we studied in intact cells [18]. Furthermore, internal perfusion may also alter signaling pathways, e.g. by washout of soluble signaling components. Finally, Takezawa and colleagues used HEK293 human embryonal kidney cells rather than the neuroblastoma cells used in this study. HEK293 cells allow substantial overexpression before they eventually die from TRPM7 expression. This allows electrophysiological characterization of channel properties with exceptional signal-to-noise ratio, but it may also cause differences in response to cellular signals. It will be interesting to include TRPM7 S1269A in such studies.

In summary, our data reveal a new level of complexity in the cAMP-dependent regulation of TRPM7, the full elucidation of which awaits further experimentation.

## Contributors

JB: Experiments. Analyzed experiments, wrote the manuscript.

ML: Conceptualized the study, performed experiments and reviewed the manuscript.

JK: Performed experiments, analyzed data and reviewed the manuscript.

YH: Performed experiments.

KJ: Conceptualized and guided the study, wrote the manuscript.

## Declaration of interest

The authors declare that they have no conflict of interest.

## Ethical standards

The authors declare that all experiments comply with Dutch and American laws.

## Acknowledgements

We thank Joachim Goedhart for providing the Twitch-2B DNA and Bram van den Broek for technical assistance.

## Funding

This work was supported by KWF grant NKI 2010-4626.

